# Engineered 3D hydrogel model reveals divergence of adhesion-migration balance in Glioblastoma under simulated microgravity

**DOI:** 10.64898/2026.04.26.720941

**Authors:** Giulia Silvani, Caitlin Williams, Nicolas Warburton, Kanwar Abhay Singh, Riddhesh Bhadresh Doshi, Yuzhu Liu, Martina Stenzel, Kate Poole, Kristopher Kilian

**Affiliations:** School of Materials Science and Engineering, UNSW Sydney, Australia; School of Chemistry, UNSW Sydney, Australia; School of Biomedical Sciences, UNSW Sydney, Australia; Australian Centre for Nanomedicine (ACN), UNSW Sydney, Australia

**Keywords:** Glioblastoma, Cancer Plasticity, Simulated Microgravity, Space mechanobiology, Cell-ECM Mechanotransduction

## Abstract

Glioblastoma is an aggressive brain cancer whose cells can switch between different modes of invasion in response to their surroundings, making the disease difficult to predict and treat. How physical forces influence this adaptability remains poorly understood. Here, we used simulated microgravity together with engineered hydrogels that independently control adhesion, degradability, and mechanical properties to test how gravity affects glioblastoma invasion. Microgravity strongly reduced invasion and shifted cells from elongated, protrusive behavior to a more cohesive state. Proteomic analysis showed reduced invasive signaling together with increased cell-matrix and cell-cell adhesion, consistent with a redistribution of contractile forces toward the cell edge. Under normal gravity, blocking CD44, integrin β1, or N-cadherin reduced matrix-dependent invasion. In contrast, under microgravity, inhibiting these same adhesion pathways restored invasion, indicating that microgravity traps cells in an overly adhesive, cohesive state that limits movement rather than motility itself. These findings show that gravity is an important regulator of cancer cell plasticity and reveal a mechanically induced vulnerability in glioblastoma invasion. More broadly, combining defined biomaterials with gravitational modulation provides a new way to study how physical forces shape tumor behavior.

## Introduction

Glioblastoma (GBM) remains one of the most lethal cancers, in large part due to its exceptional ability to adapt and invade through complex tissue environments.^1-3^ A defining feature of this plasticity is GBM’s reliance on mesenchymal invasion, a mode of motility driven by integrin-mediated adhesion, ECM remodeling, and actin-rich protrusions.^4, 5^ In this context, CD44, the principal receptor for hyaluronan, works in concert with integrins to regulate adhesion, migration, and mechanotransduction and its elevated expression has been linked to stemness and therapeutic resistance.^6-9^ Together, these interactions highlight the tumor’s deep mechanosensitivity: its capacity to sense, respond to, and exploit physical cues in its surroundings.^10, 11^ This adaptive response allows GBM cells to switch between invasion strategies, evade therapies, and resist physical or biochemical barriers^12, 13^, frequently operating through mechanisms that remain hidden under standard experimental conditions. To uncover these cryptic behaviors, we must think beyond conventional paradigms and explore unconventional environments capable of revealing how GBM reprograms its invasion machinery when fundamental physical inputs are removed or distorted.

Microgravity, defined by the near absence of gravitational force, provides a unique experimental framework to probe cellular responses under extreme mechanical disruption.^14, 15^ By eliminating gravity-driven mechanical inputs, microgravity profoundly alters human physiology, contributing to well-documented effects in astronauts such as bone loss, muscle atrophy, cardiovascular remodeling, and immune dysregulation.^16-20^ These systemic changes reflect fundamental shifts in cellular mechanosensation and signaling, suggesting that studying cells under microgravity can also uncover molecular mechanisms of adaptation relevant to disease, such as cancer.^21^ Therefore, in this context, microgravity can serve as a biophysical stress test, capable of revealing mechanosensitive vulnerabilities and compensatory mechanisms in GBM that remain invisible under normal conditions.

To explore the effects of gravitational unloading on cells, researchers commonly use ground-based simulators such as the random positioning machine (RPM). ^15, 22^ While true microgravity can only be achieved during spaceflight, access to orbital experiments is logistically challenging, costly, and limited to very few samples. By contrast, the RPM offers a rapid and reproducible approach to approximate weightlessness on Earth. These systems operate by continuously rotating biological samples along multiple axes, effectively averaging the gravity vector to near zero over time. ^23^ As a result, RPMs have become widely used tools for simulating microgravity *in vitro* and have enabled a broad range of studies in cell biology and mechanobiology.^21, 24-28^ Nevertheless, their use also introduces important limitations that must be carefully considered when interpreting results, particularly in the context of mechanosensitive behaviors. ^15, 29, 30^ Most RPM-based studies rely on standard tissue culture flasks filled with media, where cells are typically cultured in 2D or in suspension. While these approaches have been valuable for initial investigations into how cells respond to gravitational unloading, they offer limited capacity to replicate the structural and mechanical features of the extracellular matrix. As a result, key elements of cell–matrix interaction and mechanotransduction may be underrepresented. Moreover, under experimental rotation, these setups can expose cells to gradient of oxygen, uncontrolled fluid shear stress, intermittent movement, and a lack of physical matrix support, all of which may induce cellular stress responses that complicate the interpretation of microgravity-specific effects.^29^ These challenges become particularly relevant in the context of mechanosensitive tumors like GBM, whose invasive behaviour is tightly regulated by ECM parameters, e.g., composition, stiffness, topography and degradability. Without a defined matrix environment, key mechanosensitive pathways may be masked, misinterpreted, or go undetected.

To address this gap, we developed a synthetic 3D hydrogel model that enables precise decoupling of physical and biochemical variables within the tumor microenvironment. Our platform, used here for the first time in a microgravity context, is based on crosslinked hyaluronic acid (HA) and polyethylene glycol (PEG), incorporating matrix metalloproteinase (MMP)-degradable peptide sequences and RGD adhesion motifs, providing a tunable and biocompatible system to dissect GBM cell–matrix interactions in both terrestrial and microgravity contexts. We first optimized and characterized the biophysical parameters of our engineered

ECM to define conditions that modulate GBM invasion. We then applied this system to explore how GBM cells reorganize their invasion strategies when gravitational input is removed. Focusing on the adhesion network centered on CD44, and its crosstalk with integrin and N-cadherin, we uncover how microgravity triggers a selective adhesion-driven adaptation that suppresses invasion and promotes cohesive organization. Together, our findings reveal that gravitational unloading exposes a reprogrammable adhesion machinery in GBM, in which coordinated cell–ECM and cell–cell coupling emerges as the core of the adaptive response.

## Method

### Cell Culture and reagents

Human glioblastoma U87 cells were obtained from ATCC (Cat. Number HTB-4). Cells were cultured in high-glucose Dulbecco’s Modified Eagle Medium (DMEM; 4.5 g/L D-glucose, Gibco, USA) supplemented with 10% fetal bovine serum (FBS) and 1% Penicillin–Streptomycin. Cultures were maintained at 37 °C in a humidified atmosphere with 5% CO_2_, and media were replaced every 2–3 days until cells reached confluency. All experiment used U87 cells between passage numbers 25-35. For spheroid formation, GBM cells at 80–90% confluency were washed with phosphate-buffered saline (PBS; Sigma-Aldrich), detached using 0.25% Trypsin-EDTA (Gibco, Life Technologies, Cat. No. 25300054) for 3 minutes at 37 °C in 5% CO_2_, and the enzymatic reaction was neutralized by adding complete DMEM. Cell suspension was collected, centrifuged at 220 × g for 3 minutes at room temperature, and the supernatant was discarded. The resulting cell pellet was resuspended and counted. Spheroids were formed by seeding 3000 cells in 100 μL of complete DMEM into each well of a 96-well round-bottom ultra-low attachment plate (Corning), followed by incubation for 3 days at 37 °C in 5% CO_2_.

### Fabrication of PDMS Platforms for 3D Spheroid Embedding

Custom polydimethylsiloxane (PDMS) platforms were fabricated to support spheroid-laden hydrogels for both standard and simulated microgravity culture, following protocols adapted from Wang et al. (2025). ^31^ SYLGARD™ 184 Silicone Elastomer Base and Curing Agent (Dow Corning) were mixed at a 10:1 (w/w) ratio and thoroughly degassed under vacuum for at least 30 minutes to remove air bubbles. The mixture was then cast into 60 mm petri dishes and cured overnight at 60 °C. After curing, PDMS slabs were removed from the mold and cut into rectangular strips approximately 30 × 20 mm blocks. A 6 mm biopsy punch was used to create three evenly spaced wells in each PDMS platform, matching the footprint of standard 96-well plates. To assemble the platforms, PDMS surfaces and 24 × 50 mm glass coverslips were cleaned and plasma-treated for 2 minutes to activate bonding. The treated surfaces were immediately pressed together to ensure covalent attachment and stable sealing. Prior to hydrogel embedding and cell culture, bonded PDMS platforms were sterilized by exposing to UV light for 30 minutes in a biosafety cabinet.

### Synthesis of norbornene-hyaluronic acid (Nor-HA)

Norbornene-functionalized hyaluronic acid (NorHA) was synthesized following protocols adapted from *Nemec et al. (2022)*.^32^ Sodium hyaluronate (Na-HA; 40 kDa, 1013 mg, 0.0253 mmol) was first converted into its tetrabutylammonium (TBA) salt form. Na-HA was dissolved in 50 mL of Milli-Q water under vigorous stirring overnight to obtain a clear 2 wt% (20 mg/mL) solution. To perform ion exchange, 3 g of DOWEX 50WX8 resin was added and stirred for 5 hours at room temperature. The resin was subsequently removed by vacuum filtration through a 14 μm nylon mesh. The solution was then neutralized to pH 7 by the dropwise addition of approximately 300 μL of 20% TBA hydroxide (TBA-OH). The neutralized HA-TBA solution was aliquoted into 20 mL portions, frozen at –80 °C, and lyophilized to yield a light pink, low-density solid. For functionalization, the lyophilized HA-TBA was dissolved in anhydrous dimethyl sulfoxide (DMSO) to a final concentration of 2 wt% under an inert argon atmosphere and stirred vigorously for 2 hours. To this solution, 25 equivalents of 5-norbornene-2-methylamine, 30 equivalents of PyBOP, and 30 equivalents of DIPEA were added to target a degree of functionalization (DOF) of 25%. After 2 hours of reaction, the mixture was neutralized by the addition of cold water and transferred into a dialysis membrane (molecular weight cutoff: 6–8 kDa). Dialysis was performed against 5 g/L NaCl for 3 days, followed by Milli-Q water for an additional 3 days, with dialysate exchanged twice daily throughout. The final dialyzed solution was lyophilized to yield a white, low-density NorHA powder, which was stored at –20 °C. The degree of functionalization was confirmed by proton nuclear magnetic resonance (^1H NMR, 400.17 MHz, D_2_O).

### Synthesis of functionalized PEG Nor-HA and rheology test

Nor-HA was dissolved in basal DMEM with stirring at room temperature overnight to yield a final concentration of 2.0% w/w (0.5 mM). 4-arm polyethylene glycol thiol (10 kDa) was dissolved in deionized (DI) water to yield a final concentration of 1% w/w (1.0 mM). GRGDSC and GRDGSC peptides were dissolved in DI water after 20-minute UV-sterilization to yield a final concentration of 100 mM. Matrix metalloproteinase (MMP) degradable (Deg; GCGPQGIWQCG) and non-degradable (non Deg; GCGpQGIWGQCG) cross-link sequences were UV-sterilised for 20 minutes and dissolved in DMSO for a final concentration of 100 mM. Lithium phenyl-2,4,6-trimethylbenzoylphosphinate (LAP) was dissolved in water at 2.5% w/w (85 mM) and stored in foil between experiments at 4°C. PEG, NorHA and DI water were filtered with 0.22 *μ*m membrane filters to sterilise prior to any cell work. 200 *μ*L gel stock solutions were prepared via addition of 100 *μ*L 2% NorHA (1%, 0.25 mM final), 16.48/32.95/49.43 *μ*L of 1% PEG (5, 10 and 15% functionalisation of norbornene respectively; 1.65 × 1 ^−8^, 3.30 × 10^−8^ and 4.94 × 10^−8^ moles respectively), 5.93 *μ*L (5.93× 10^−7^ mol) RGD/RDG, 2.64 *μ*L (2.64× 10^−7^ mol) MMP+/- and 4.00 *μ*L 2.5% LAP. The total volume was made up to 200 *μ*L with DI water. Rheological measurement of the PEG-NorHA gels were conducted using an Anton Paar MCR 302 rheometer fitted with a 25 mm diameter parallel plate. For each measurement, 600 *μ*l of gel stock solution was placed between the plates with a 1mm gap. A time sweep test was performed applying 0.2% strain at a frequency of 1 Hz. UV light was shone onto the sample through the glass stage for 3 minutes while the time sweep data was collected. The storage modulus (G′) was extracted from the time sweep curves and analysed in GraphPad Prism. The final plateau value of G′ was used as the representative elastic modulus for each hydrogel formulation once crosslinking reached equilibrium.

### Spheroid culture into engineered hydrogels

GBM spheroids, PDMS platforms, and PEG-NorHA hydrogel stock solutions were prepared as described in Sections above. To begin the embedding process, 30 *μ*L of hydrogel stock solution was dispensed into each PDMS well, ensuring uniform and level coverage at the base and expose to UV light for 3 minutes to crosslink. Spheroids were prepared by carefully removing the culture media from each well using a pipette. A 20 *μ*L aliquot of fresh hydrogel stock was then added to each spheroid well. Each spheroid was immediately collected along with the hydrogel mixture using a pipette and transferred onto the pre-formed hydrogel base within the PDMS wells. This process was repeated for all spheroids within a given experimental condition. Following placement, PEG-NorHA hydrogel was again crosslinked using UV light for 3 minutes. Once all spheroids were embedded, the hydrogels were incubated for ∼10 minutes at 37 °C to ensure full gelation. Subsequently, 150 *μ*L of complete DMEM was gently added on top of each well and the platform immediately used for simulated microgravity experiments.

### Simulated microgravity experiments

Microgravity was simulated using a Random Positioning Machine (RPM) (EXPLOR Space Technologies, Sydney, NSW, Australia). Immediately before placement in the RPM, each platform was filled with media and then sealed with an air-permeable adhesive membrane. This was done to exclude air bubbles and avoid leaking during the rotation of the sample mounted in the microgravity simulator. Subsequently, samples were mounted on the RPM at the center of the inner frame to ensure simulation of microgravity (∼0.001 g), while eliminating artefacts arising from rotational kinematics. The RPM was located inside a cell culture incubator set for standard cell culture conditions, and a duplicate sample placed in the same incubator, next to the RPM, as a 1g control. Simulated microgravity was then applied using standard, validated factory settings within the EXPLOR software for 24 hours. Post-application of simulated microgravity, platforms were removed from the incubator and cells were either collected for further analysis or fixed overnight at 4 °C, washed, and stored in phosphate-buffered saline (PBS) protected from light at 4 °C until further processing.

### Inhibition Treatment

To investigate the functional contribution of adhesion-associated receptors to GBM invasion under normal gravity and simulated microgravity, pharmacological inhibition of β1-integrin, αvβ3-integrin, CD44, and N-cadherin was performed. U87 spheroids were generated as described above and embedded within adhesive and degradable NorHA–PEG hydrogels. For integrin inhibition, function-blocking antibodies targeting β1-integrin (α5β1-specific; Merck, Cat. MAB2514) or αvβ3-integrin (Merck, Cat. MAB1976) were incorporated directly into the hydrogel precursor solution prior to crosslinking at a final concentration of 10*μ*M. For CD44 inhibition, a blocking antibody against CD44 (Invitrogen, Cat. MA5-13890) was used at 30 µg/mL. To block CD44 receptors on the surface of the cells, spheroids were incubated for 15 minutes in media supplemented with CD44 antibody. Spheroids were then encapsulated in hydrogel solution as described above. N-cadherin inhibition was performed using a function-blocking antibody (Abcam, Cat. D4R1H) at 10 µg/mL. Immediately following hydrogel encapsulation and prior to simulated microgravity exposure, inhibitor-containing media was gently added on top of each PDMS well to ensure continuous exposure throughout the 24-hour culture period. Samples were then either mounted in the Random Positioning Machine (RPM) to simulate microgravity (∼0.001 g) or maintained under standard 1g incubator conditions for 24 hours, as described in the “Simulated microgravity experiments” section. Following treatment, spheroids were either fixed for immunofluorescence analysis or processed for downstream assays, including invasion quantification and CD44 expression analysis.

### Immunofluorescence

Following fixation and PBS storage, 3D hydrogel-embedded spheroids were permeabilized using 0.1% Triton X-100 in PBS for an overnight at 4 °C. After permeabilization, 3D hydrogel-embedded spheroids were washed in PBS for 10 minutes and primary antibodies were diluted 1:200 in PBS and 50 *μ*L was added to each well. Samples were incubated overnight at 4 °C in the dark. After primary antibodies, 3D hydrogel-embedded spheroids were washed in PBS for 10 minutes again and secondary antibodies were diluted 1:200 in PBS and 50 *μ*L was added to each well. Samples were incubated overnight again at 4 °C in the dark, washed the day after in PBS for 10 minutes and stored at 4 °C in the dark until imaging. For this study, mouse anti-CD44 (1:200 dilution; Invitrogen, Cat. MA5-13890); mouse anti-Nestin (1:200, Invitrogen, Cat. MA5-13890); rabbit anti-Ki67 (Sigma, Cat. SAB5600249); rabbit anti-pMLC (CellSignaling, Cat. 3671S) and rabbit anti-N-cadherin (1:200 dilution; Abcam, Cat. D4R1H) were used to label respective targets. Fluorescent secondary antibodies were anti-mouse Alexa Fluor 488 (1:200, Abcam) and anti-rabbit Alexa Fluor 555 (1:200, Abcam). Actin cytoskeleton was stained with Alexa Fluor 647-Phalloidin (1:200, ThermoFisher, Cat. Number A30107) and nuclei were stained with Hoechst (0.1 *μ*g/mL in PBS, ThermoFisher, Cat. H3570).

### Imaging and analysis

Samples were imaged using the Leica TCS SP8 Digital LightSheet (DLS) and Olympus FV4000 both equipped with the 10x air objective lens to collect image stacks encompassing the entire spheroid and surrounding dissociated cells. Light intensity settings for all markers were kept consistent across samples, with DAPI settings varied to optimize visualization of the cell nuclei. Spheroids were analyzed using Imaris 9.1.2 software to segment cells by creating masks with a cut-off filter of 200 µm^3^ to exclude cell debris. Dissociated cells were filtered by center of image mass versus z depth, where the z value was determined by orientating the 3D image stacks to the plane of view showing the cells on the glass. A spheroid mask corresponding to spheroid size at time = 0 was created and dissociated cells outside this masked area were identified in Imaris. From these data, the number of dissociated cells, their coordinate positions, and fluorescent intensity on different channels were exported and analyzed. Immunofluorescence images of GBM spheroids stained with phalloidin were analyzed in ImageJ (Fiji) to assess cytoskeletal protrusion morphology. Images were first converted to 8-bit and thresholder to generate binary masks, which were then processed using the Skeletonize tool to reduce protrusive actin structures to one-pixel-wide lines while preserving their topology. The resulting skeletons were quantified with the Analyse Skeleton (2D/3D) plugin, which automatically provides morphometric parameters such as the number of branches and maximum branch length from the dialog output. These values were used to evaluate differences in protrusive organization and cytoskeletal network complexity across experimental conditions.

### Protein quantification of cell spheroids and cell secretome

Conditioned medium was collected from each group after 24 hours of spheroid culture in NorHA-PEG gels, centrifuged at 3000xg for 5 minutes, and subsequently utilized for secretome profile analysis. Hydrogels from both groups were then collected from the PDMS platform and digested using RASTRUM™ Cell Retrieval Solution (Inventia, Catalog no. F235). The solution with spheroids was centrifuged at 3000xg for 5 minutes and cells were rinsed three times with PBS. Lysis was performed using 100 µL RIPA buffer (Life Technologies 89900) in the presence of a protease inhibitor cocktail (Sigma 11836153001), with samples vortexed at 4 °C every minute. The cell lysates were then centrifuged at 3000xg for 5 minutes to eliminate residual gel debris, and the supernatant was collected for subsequent profile analysis. Protein concentrations were determined using the Pierce BCA Protein Assay Kit.

### Proteomic analysis of cell secretome and cell spheroid lysates

The secretomic samples were initially concentrated using a 3 kDa spin cartridge, followed by protein denaturation with 10 M urea. The denatured proteins were then reduced (5 mM DTT, 37 °C, 30 min), alkylated (10mM IA, RT, 30 min), and digested with trypsin overnight. The lysate samples were prepared with the same protocol without concentration and denaturation steps. Samples were desalted with two SDB-RPS disks (Empore, Sigma Cat#66886-U) packed in a 200 *μ*L pipette tip as described previously. Extracted peptides from each clean-up were reconstituted in 10 *μ*L 0.1% (v/v) formic acid, 0.05% (v/v) heptafluorobutyric acid and 2% acetonitrile in water ^33^. Digested peptides were separated by nanoLC using an Ultimate nanoRSLC UPLC and autosampler system (Dionex, Amsterdam, Netherlands). Samples (2.5 µl) were concentrated and desalted onto a micro C18 precolumn (300 µm x 5 mm, Dionex) with H2O:CH3CN (98:2, 0.1 % TFA) at 15 µl/min. After a 4 min wash the pre-column was switched (Valco 10 port UPLC valve, Valco, Houston, TX) into line with a fritless nano column (75µ x ∼30 cm) containing C18AQ media (1.9µ, 120 Å Dr Maisch, Ammerbuch-Entringen Germany). Peptides were eluted using a linear gradient of H2O:CH3CN (98:2, 0.1 % formic acid) to H2O:CH3CN (64:36, 0.1 % formic acid) at 200 nL/min over 30 min and 90 min for supernatant samples and lysate samples, respectively. High voltage 2000 V was applied to low volume Titanium union (Valco) and the tip positioned ∼0.5 cm from the heated capillary (T=275°C) of a Orbitrap Fusion Lumos (Thermo Electron, Bremen, Germany) mass spectrometer. Positive ions were generated by electrospray and the Fusion Lumos operated in data dependent acquisition mode (DDA). A survey scan m/z 350-1750 was acquired in the orbitrap (resolution = 120,000 at m/z 200, with an accumulation target value of 400,000 ions) and lock mass enabled (m/z 445.12003). Data-dependent tandem MS analysis was performed using a top-speed approach (cycle time of 2s). MS2 spectra were fragmented by HCD (NCE=30) activation mode, and the ion-trap was selected as the mass analyser. The intensity threshold for fragmentation was set to 25,000. A dynamic exclusion of 20 s was applied with a mass tolerance of 10ppm. Peak lists were generated using Mascot Daemon/Mascot Distiller (Matrix Science, London, England) or Proteome Discoverer (Thermo, v1.4) using default parameters and submitted to the database search program Mascot (version 3.1, Matrix Science). Search parameters were Precursor tolerance 4 ppm and product ion tolerances ± 0.5 Da; Met (O) carboxyamidomethyl-Cys specified as variable modification, enzyme specificity was trypsin, 1 missed cleavage was possible and the Uniprot (January 2024) was searched.

### Extracellular Vesicle isolation and quantification

Conditioned media was collected after 24 hours of cell culture and centrifuged at 3500×g for 5 minutes to remove cells, followed by filtration through a 0.22 *μ*m sterile filter to eliminate debris. Extracellular vesicles (EVs) were isolated using the Total Exosome Isolation Reagent (ThermoFisher Scientific, Cat: 4478359) by mixing the conditioned media with the reagent in a 1:0.5 ratio and incubating overnight at 4°C. The mixture was then centrifuged at 10000×g for 1 hour at 4°C, and the resulting EV pellet was resuspended in 100 *μ*L of 4% trehalose solution in PBS. The final EV preparation was stored at −80°C until further use. Extracellular vesicle (EV) samples were thawed, vortexed, and diluted with 1× sterile PBS in 1:100 ratio prior to analysis. Measurements were performed using the manufacturer’s default software settings for EVs, scanning 11 cell positions with a camera sensitivity of 65, shutter speed of 200, and cell temperature maintained at 25°C. Videos captured during the scan were analysed using Zetaview Software version 8.02.31, applying specific parameters including a maximum particle size of 1000, minimum particle size of 5, and an embedded laser wavelength of 405 nm.

### Tumorsphere assay

To assess the self-renewal and stem-like properties of U87 under normal and microgravity condition, we employed a tumorsphere assay using ultra-low attachment culture plates following protocols adapted from *Silvani et al*. (2025).^34^ U87 cells were cultured in normal and microgravity conditions in dedicated platform previously developed for microgravity experiments ^35^. After simulated microgravity experiment, cells were detached using 0.25% Trypsin-EDTA (Gibco, Life Technologies, Cat. No. 25300054) for 3 minutes at 37 °C in 5% CO_2_, and the enzymatic reaction was neutralized by adding complete DMEM. Cell suspension was collected, centrifuged at 220 × g for 3 minutes at room temperature, and the supernatant was discarded. The resulting cell pellet was resuspended and counted. Cells were then seeded in 96-well ultra-low attachment plates at a concentration of approximately 3 × 10^3^ cells per well into the serum-free media (R&D Systems™ StemXVivo Serum-Free Tumorsphere Media, Cat. Number CCM012). The plates were incubated at 37 °C with 5% CO_2_, and tumorsphere formation was monitored over 7 days, changing media each 48 hours. Tumorsphere were observed using bright-field microscopy and the size of spheres was quantified using ImageJ software.

### Statistical analyses

Unless otherwise noted, all experimental results are from at least three independent experiments. Error bars represent the standard error of the mean (SEM). For comparisons between two groups, a two-sided unpaired t-test was used. For comparisons involving two factors, two-way ANOVA with Tukey’s post hoc test was used. All analyses were performed using GraphPad Prism v8.2.0 (GraphPad Software). Statistical details are provided in the figure legends. P-value is reported for statistical significance. Comparisons between samples were considered to be statistically significant if the p-value was *p < 0.05, **p < 0.01, ***p < 0.001, ***p < 0.0001.

## Result

### A tunable NorHA-based hydrogel system recapitulates ECM-driven invasion sensitivity in GBM

To establish a reductionist yet brain-relevant ECM mimic, we designed a modular system employing synthetic norbornene-functionalized hyaluronic acid (NorHA) that is amenable to crosslinking via thiol-ene click chemistry **(Fig. 1A)**. Successful modification was confirmed by ^1^H NMR, which showed characteristic norbornene peaks at δ = 5.8–6.3 ppm in addition to the native HA signals at δ = 2.9–3.9 ppm **(Fig. S1)**. Degree of functionalization (DOF) was quantified by peak integration, yielding ∼35% substitution of the HA backbone. This level of substitution was consistent across batches and provided sufficient reactive groups for thiol–ene crosslinking while preserving the biochemical relevance of HA. Using thiol–ene chemistry, we introduced either integrin-binding R<u>G</u>D peptides or their scrambled control R<u>D</u>G, together with either MMP-degradable or non-degradable crosslinkers, thus independently controlling matrix adhesiveness and degradability **(Fig. 1B)**. To test how these parameters influence tumor behavior, pre-formed U87 spheroids were embedded within the matrices using a sandwich-style configuration, ensuring complete three-dimensional encapsulation **(Fig. 1C)**. After 24 hours of culture, immunofluorescence staining with phalloidin revealed robust outgrowth in adhesive (R<u>G</u>D), degradable conditions (Deg), with cells extending elongated, actin-rich protrusions into the surrounding matrix. In contrast, spheroids in non-adhesive (R<u>D</u>G) or non-degradable (Non Deg) gels remained compact and rounded, with minimal dissemination **(Fig. 1D)**. To quantify these observations, we measured the number of invading cells that migrated away from the spheroid core across conditions. Consistent with the imaging, invasion was significantly enhanced in adhesive and degradable matrices (R<u>G</u>D-Deg) but strongly suppressed when either adhesion or degradability was absent **(Fig. 1E)**. Together, these results confirm that the platform reliably recapitulates ECM-dependent regulation of GBM invasion, establishing adhesion and degradability as the primary drivers of invasive behavior in our system.

**Fig. 1.**
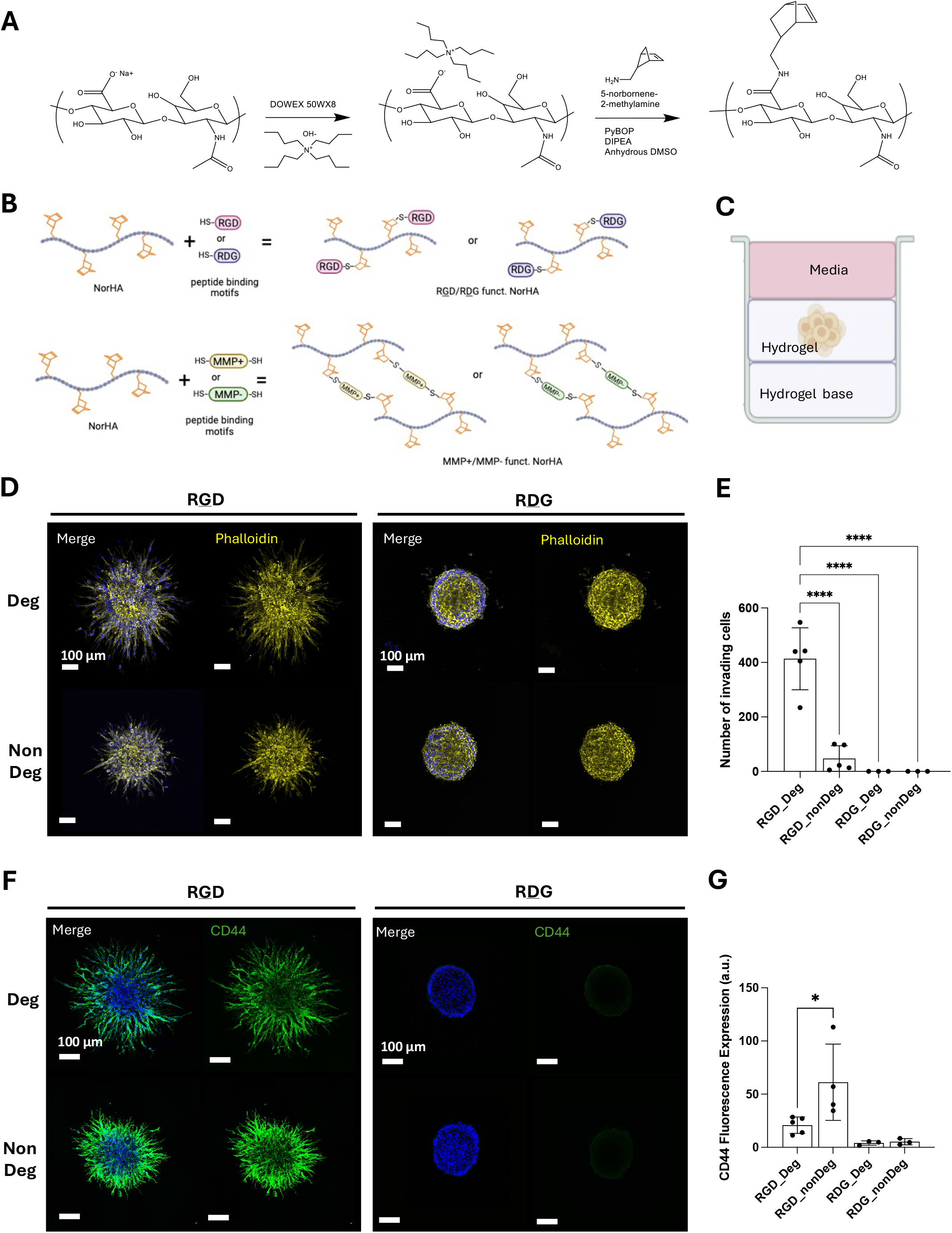
Characterization of ECM adhesiveness and degradability. **(A)** Schematic of the modular norbornene-functionalised hyaluronic acid (NorHA) hydrogel platform engineered using thiol–ene click chemistry to enable independent tuning of matrix adhesiveness and degradability. **(B)** Schematic showing that NorHA macromers were functionalised with either integrin-binding R<u>G</u>D peptides or scrambled R<u>D</u>G controls and crosslinked using either matrix metalloproteinase (MMP)-cleavable or non-cleavable peptides, generating four distinct ECM conditions. **(C)** Experimental configuration for three-dimensional tumour culture: pre-formed GBM spheroids were placed between two hydrogel layers to ensure complete encapsulation and uniform matrix contact on all spheroids sides. **(D)** Representative immunofluorescence images of spheroids after 24 h culture in the different matrix conditions, stained with phalloidin (F-actin, yellow) and DAPI (nuclei, blue). Scale bars: 100 *μ*m. **(E)** Quantification of invading cells across matrix conditions expressed as the mean ± SEM from five independent experiments, except for RDG_Deg and RDG_nonDeg (n=3). **(F)** Representative immunofluorescence images showing CD44 immunofluorescence (green) and DAPI (blue) in U87 spheroids cultured under the indicated conditions. Scale bars: 100 *μ*m. **(G)** Quantification of CD44 fluorescence intensity expressed as the mean ± SEM from five independent experiments, except for RDG_Deg and RDG_nonDeg (n=3).

We next asked whether the same matrix parameters also influence cell plasticity. To gauge changes in phenotype, we focused on CD44, a principal hyaluronan receptor implicated in GBM invasion and stem cell-like characteristics. Immunofluorescence analysis revealed that CD44 expression was strongly upregulated in non-degradable matrices (R<u>G</u>D-NonDeg), whereas expression was lower in degradable gels (R<u>G</u>D-Deg) despite robust invasion **(Fig. 1F)**. Additionally, CD44 levels were negligible in non-adhesive controls (R<u>D</u>G), suggesting that its induction requires integrin engagement. Quantification of fluorescence intensity within spheroids demonstrated that the highest CD44 expression occurred in adhesive but non-degradable conditions **(Fig. 1G)**, suggesting that adhesion under confinement influences cell plasticity.

To modulate ECM mechanics while keeping biochemical signals constant, we crosslinked NorHA with bioinert 4-arm PEG-thiols **(Fig. 2A)**, varying the degree of Nor:SH ratio (80:1, 40:1, 27:1, corresponding to 5%, 10% and 15% respectively) to tune network density and stiffness. Importantly, the concentration of adhesion and degradation peptide motifs was kept constant across formulations. Mechanical testing confirmed a PEG-dependent increase in storage modulus, ranging from ∼ 90 to 190 Pa, corresponding to the soft brain-like ECM range commonly used to model healthy brain parenchyma conditions (**Fig. 2B, C)**. Because stiffness in these hydrogels is modulated through polymer concentration, this mechanical tuning is inherently coupled to changes in network architecture, including mesh size and molecular crowding, which together define the physical microenvironment experienced by the cells. After culture for 24 hours, immunofluorescence staining of actin revealed that cell invasion decreased progressively across this mechanical range, even in degradable matrices, with spheroids in stiffer gels showing markedly reduced dissemination compared to softer counterparts **(Fig. 2D)**. Invasion was further impaired in non-degradable conditions, where spheroids remained compact and largely confined within the gel. Quantification of disseminated cells confirmed that matrix stiffness reduced invasion regardless of degradability, underscoring that both mechanical confinement and matrix remodelling are critical regulators of GBM motility **(Fig. 2E)**. Interestingly, Ki67 staining **(Fig. 2F)** revealed a parallel effect on proliferation: spheroids in stiffer matrices displayed markedly higher proliferative indices **(Fig. 2G)**, consistent with stiffness-driven activation of mechanically regulated growth pathways. To further dissect this effect, we next examined whether matrix stiffness also modulates CD44 expression. Embedding spheroids into R<u>G</u>D-functionalised hydrogels with increasing PEG content revealed a progressive rise in CD44 intensity, with the strongest signal observed in the stiffest gels **(Fig. 2H)**. When degradability was simultaneously restricted, CD44 levels were further elevated, indicating that stiffening and non-degradability act as complementary triggers of receptor upregulation **(Fig. 2I)**.

**Fig. 2.**
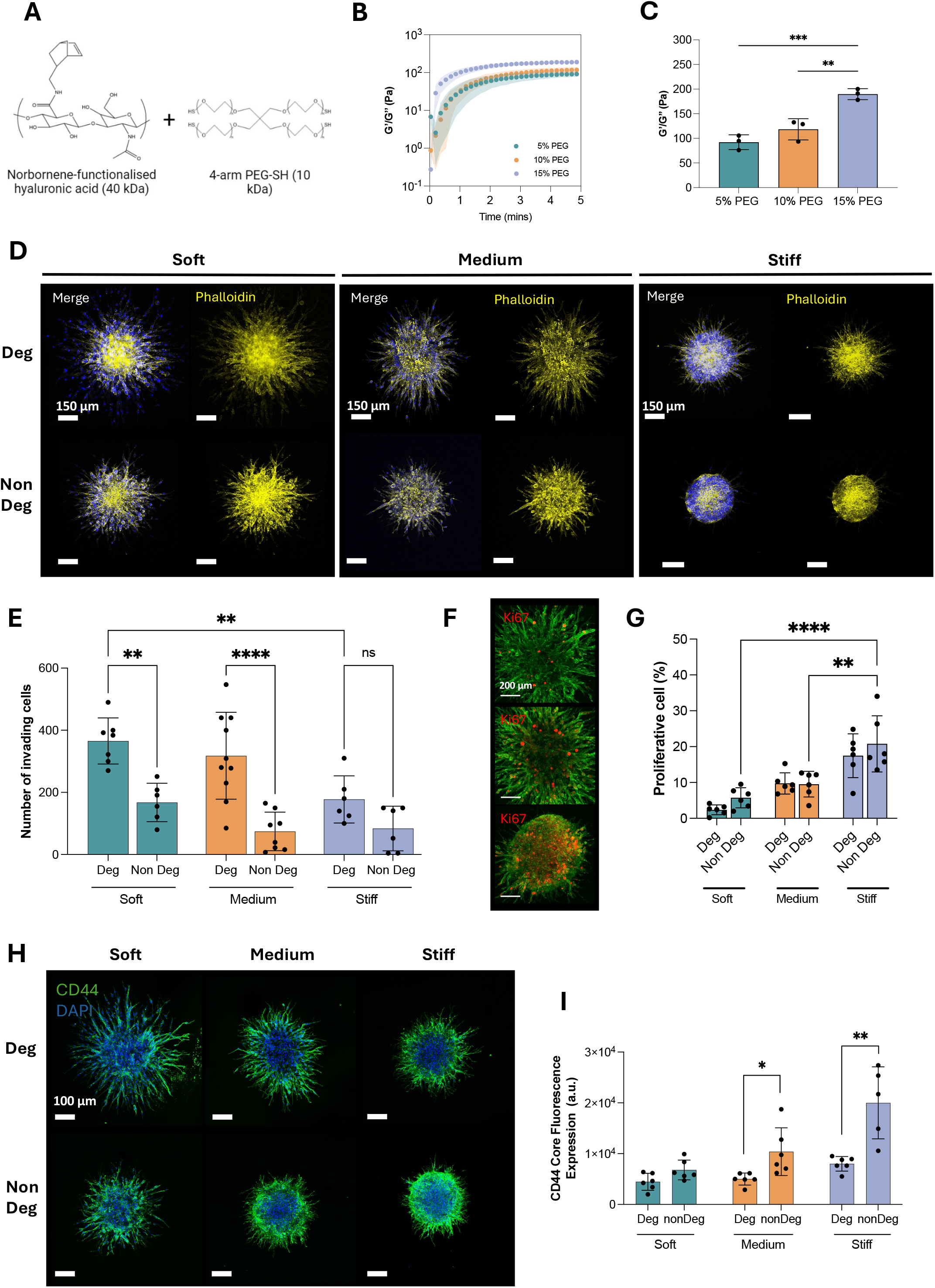
Characterization of matrix stiffness in the engineered ECM and corresponding GBM spheroid response. **(A)** Schematic of the NorHA–PEG hydrogel formulation. NorHA was crosslinked with 4-arm PEG-SH to tune network density and stiffness. **(B)** Representative time sweep of storage (G′) and loss (G″) moduli during in situ photopolymerization, showing rapid gelation and stabilisation of mechanical properties over time for hydrogels formulated with increasing PEG content (5%, 10%, 15%). **(C)** Quantification of the plateau storage modulus (G′) following gelation, demonstrating stiffness tuning across formulations. Values correspond to the mean ± SD of independent measurements (n = 3), with the final G′ value extracted after stabilisation of the time sweep. **(D)** Representative immunofluorescence images of U87 spheroids after 24 h culture in adhesive matrices of increasing stiffness, stained with phalloidin (F-actin, yellow) and DAPI (nuclei, blue). Scale bars: 150 μm. **(E)** Quantification of disseminated cells migrating across stiffness and degradability conditions, expressed as the mean ± SEM from at least 6 independent experiments. **(F)** Representative immunofluorescence images of Ki67 immunostaining (red) showing increased proliferative activity in spheroids cultured within stiffer matrices. Scale bar: 200 μm. **(G)** Quantification of Ki67-positive cells (proliferative index) across stiffness and degradability conditions expressed as the mean ± SEM from 6 independent experiments. **(H)** Representative immunofluorescence images of CD44 immunofluorescence (green) and DAPI (blue) in spheroids embedded in adhesive matrices of increasing stiffness. Scale bars: 100 μm. **(I)** Quantification of CD44 fluorescence intensity across stiffness and degradability conditions expressed as the mean ± SEM from 6 independent experiments, except for Stiff_nonDeg (n=5).

Together, these findings demonstrate that our NorHA-based hydrogel system provides a tunable environment to model GBM. Beyond reproducing canonical ECM-dependent invasion, the platform reveals coordinated changes in invasion morphology and CD44 expression in response to defined adhesive and mechanical constraints. This establishes a robust baseline from which to investigate how GBM cells reorganize their invasion strategies under the unconventional mechanical stress of microgravity.

### Mechanical unloading suppresses mesenchymal invasion in favor of adhesion-/cohesion-driven adaptation

To evaluate the effects of simulated microgravity on GBM invasion traits, we selected the intermediate stiffness condition (10%-PEG/NorHA gels). U87 spheroids were embedded and cultured either in the Random Positioning Machine (RPM) for 24 hours or in parallel control samples kept in the incubator under standard 1g conditions. Under simulated microgravity exposure, immunofluorescence staining with phalloidin and DAPI revealed a marked reduction in cell dissemination from spheroids compared to 1g controls, but this effect was specific to adhesive and degradable matrices (R<u>G</u>D Deg) **(Fig. 3A, B)**. In non-adhesive (R<u>D</u>G) and non-degradable (Non Deg) gels, where invasion was already impaired due to the absence of cell–matrix anchoring and degradable crosslinks, microgravity did not further suppress invasion **(Fig. 3B)**. This result suggests that the inhibitory effect of simulated microgravity requires an active invasion program, and that mechanical unloading interferes with integrin-and protease-dependent migration. In parallel, Ki67 immunostaining of spheroids grown in adhesive and degradable environment, demonstrated reduced cell proliferation under simulated microgravity, as shown in **Fig. S2A** and quantified in **Fig. S2B**. Strikingly, analysis of spheroid Z-stacks showed a switch in invasion mode. While under normal gravity cells disseminated individually with elongated, mesenchymal protrusions; under simulated microgravity cells invaded as rounded clusters, maintaining cell–cell contacts **(Fig. 3C)**. To quantify this change, we measured the length of protrusions and the number of branches extending from the spheroid using skeletonization analysis **(Fig. 3D, F)**. Both metrics were significantly reduced under simulated microgravity condition, confirming that the protrusive network was compressed and less elaborated compared to 1g controls. These observations suggests that gravitational unloading shifts the balance from a mesenchymal to a collective/amoeboid-like invasive phenotype.

**Fig. 3.**
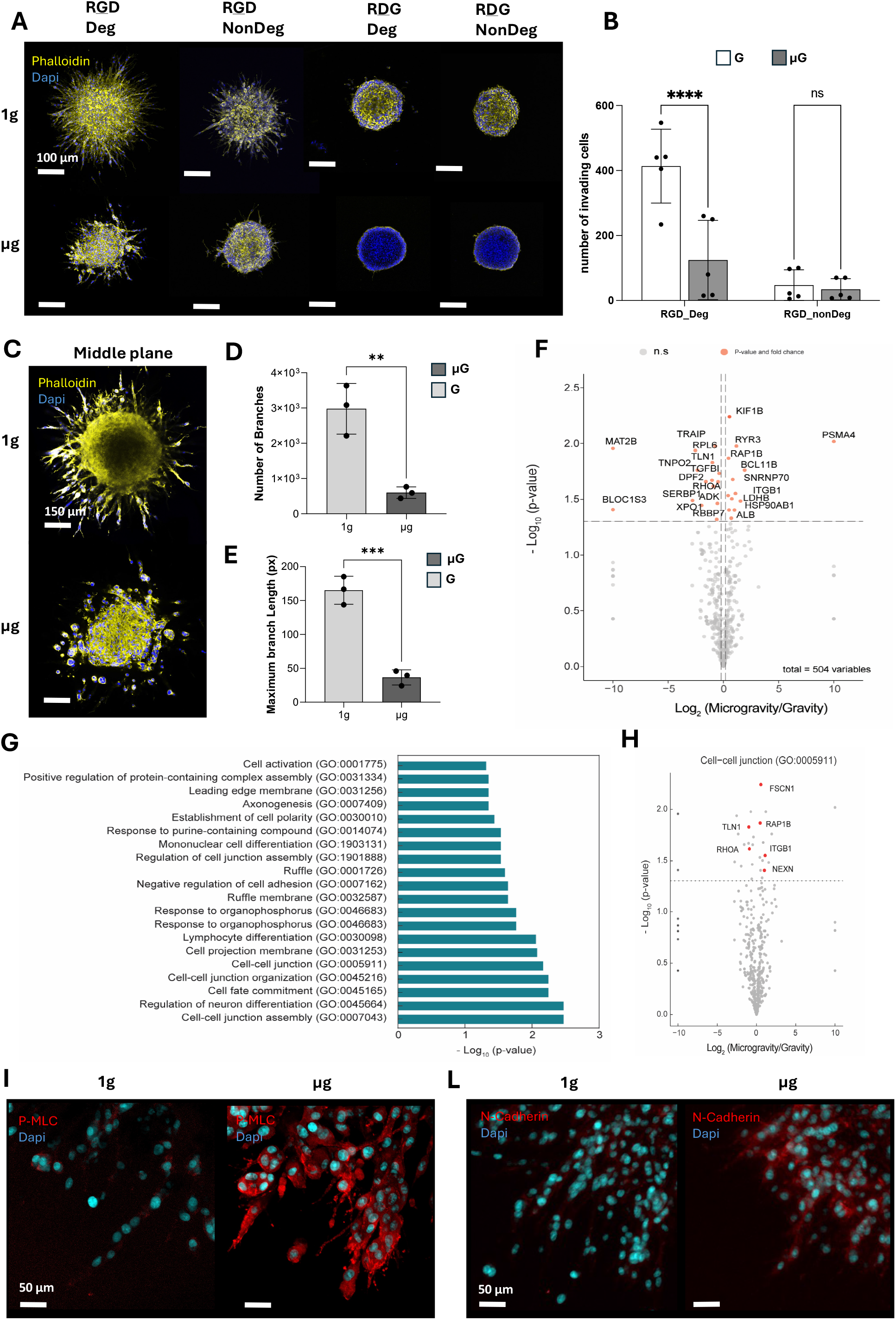
Characterization of GBM spheroid invasion, morphology and proteomic response under simulated microgravity. **(A)** Representative immunofluorescence images of U87 spheroids embedded in adhesive NorHA hydrogels (medium stiffness) and cultured for 24 h under normal gravity (1g) or simulated microgravity (μg). Spheroids were stained with phalloidin (F-actin, yellow) and DAPI (nuclei, blue). Scale bars: 100 μm. **(B)** Quantification of disseminated cells across matrix conditions under 1g and s-μg, expressed as the mean ± SEM from 5 independent experiments. **(C)** Middle plane of the reconstructions (Z-stacks) illustrating invasion mode under 1g and simulated microgravity (μg). Scale bars: 100 μm. Quantification of the number of branches elongated from the spheroid core **(D)** and the maximum branches length obtained with skeletonization analysis under 1g and μg conditions **(E)**, both expressed as the mean ± SEM from 3 independent experiments. **(F)** Volcano plot showing differentially expressed proteins in cell lysates from spheroids cultured under μg compared to 1g (log_2_ fold change vs −log_10_ p-value). **(G)** Gene Ontology (GO) enrichment analysis of differentially expressed proteins, revealing enrichment of processes related to cell–cell junction assembly and regulation of cell adhesion. (**H**) Volcano plot showing differentially expressed proteins for cell-cell junction (GO:0005911). **(I)** Representative immunofluorescence images of phosphorylated myosin light chain (pMLC, red), phalloidin (yellow), and DAPI (blue), showing minimal pMLC under 1g and a pronounced cortical pMLC distribution under μg. Scale bars: 50 μm. **(L)** Representative immunofluorescence images of N-cadherin immunostaining (red), phalloidin (yellow), and DAPI (blue). Scale bars: 100 μm.

To probe the molecular basis of this hypothesis, we performed a proteomic profiling of GBM spheroids (cell culture media and cell lysate) cultured under conditions of gravity and simulated microgravity. Culture media collected from spheroids grown under microgravity conditions showed an upregulation of proteins involved in the process of vesicle formation **(Fig. S3A)**, specifically AOC3 (Log_2_ (fold change) = 2.23), GC (Log_2_ (fold change) = 0.40) and AHSG (Log_2_ (fold change) = 0.56). Direct EVs quantification, using particle tracking analyses, further confirmed this result, showing a ∼1.25-fold increase in vesicle release under simulated microgravity compared to 1g controls **(Fig. S3B)**, with size distribution remaining within the small EV range (30–150 nm) **(Fig. S3C, D)**. This result suggests that mechanical unloading increases vesicle output rather than altering subtype. Similarly, analysis of the cell lysate shows differential expression of 30 proteins in spheroids cultured under simulated microgravity conditions **(Fig 3F)**. To identify the broader cellular mechanisms associated with these differentially expressed proteins, we utilized Gene Ontology (GO) enrichment analysis **(Figure 3G)**. GO analysis indicates enrichment of cellular processes such as cell-cell junction assembly (GO:0007043) and negative regulation of cell adhesion (GO:0001726) respectively. Specifically looking at processes such as cell-cell junctions, we observe an upregulation of the associated proteins NEXN, ITGB1, FSCN1 and RAP1B, and downregulation of RHOA, a master regulator in cell motility **(Figure 3H)**. The proteomic profile supports the imaging data, indicating a shift toward a hyper-adhesive state characterized by concurrent strengthening of cell–cell junctions and cell–matrix anchoring under simulated microgravity conditions.

To probe whether these altered adhesive phenotypes correspond with changes in actomyosin contractility, we immunostained for phosphorylated myosin light chain. Immunofluorescence analysis revealed a striking difference between gravity and simulated microgravity conditions **(Figure 3I)**. Under conditions of normal gravity, pMLC staining was minimal or absent, consistent with the predominance of a mesenchymal invasion mode driven by focal adhesion and cytoskeletal tension. By contrast, GBM cultured under microgravity demonstrated a distinct cortical distribution of pMLC2 around the cell periphery, a hallmark of amoeboid-like, contractility-driven motility. To support the enrichment of cell–cell junction–related processes identified in the proteomic analysis, we next immunostained for the cell–cell adhesion molecule N-cadherin. While overall N-cadherin intensity remained low, spheroids exposed to simulated microgravity displayed a more distinct and localized junctional signal, indicating enhanced cell–cell engagement and consistent with a shift toward adhesion-dependent multicellular organization **(Figure 3L)**.

### Mechanical unloading redirects GBM adaptation toward HA-guided CD44 adhesion rather than stemness

Having identified CD44 as a receptor selectively upregulated when invasion is restricted in our model matrices, we next asked whether mechanical unloading in simulated microgravity would influence this adaptive response. Immunofluorescence analysis revealed that, despite the overall suppression of invasion, CD44 expression was significantly elevated in microgravity compared to 1g controls **(Fig. 4A)**. This effect was most pronounced in adhesive and degradable matrices (R<u>G</u>D Deg), where invasion was otherwise permitted under normal gravity but became strongly reduced in microgravity **(Fig. 4B)**. In non-adhesive (R<u>D</u>G) and non-degradable (Non Deg) matrices, where invasion was already constrained, CD44 expression did not follow a consistent trend across samples **(Fig. 4B, C)**, consistent with limited engagement of adaptive invasion-associated responses. To determine whether CD44 upregulation under microgravity depended specifically on hyaluronan engagement, we assessed GBM spheroids embedded in alginate hydrogels, which provide a mechanically comparable but CD44-nonbinding matrix. However, to control for potential effects of cell–matrix adhesion, alginate hydrogels were tested in both non-adhesive and adhesive (RGD-modified) formulations. Immunofluorescence analysis revealed comparable CD44 staining patterns across conditions, and simulated microgravity did not induce a detectable increase in CD44 expression in either adhesive or non-adhesive alginate matrices **(Fig. S4A)**. Quantification of CD44 signal intensity confirmed the absence of significant differences between gravity conditions or between adhesive states **(Fig. S4B)**. Together, these results indicate that CD44 upregulation under simulated microgravity requires engagement with the HA backbone and cannot be recapitulated by integrin-mediated adhesion alone, suggesting that HA–CD44 interactions play a dominant role in driving this adaptive response.

**Fig. 4.**
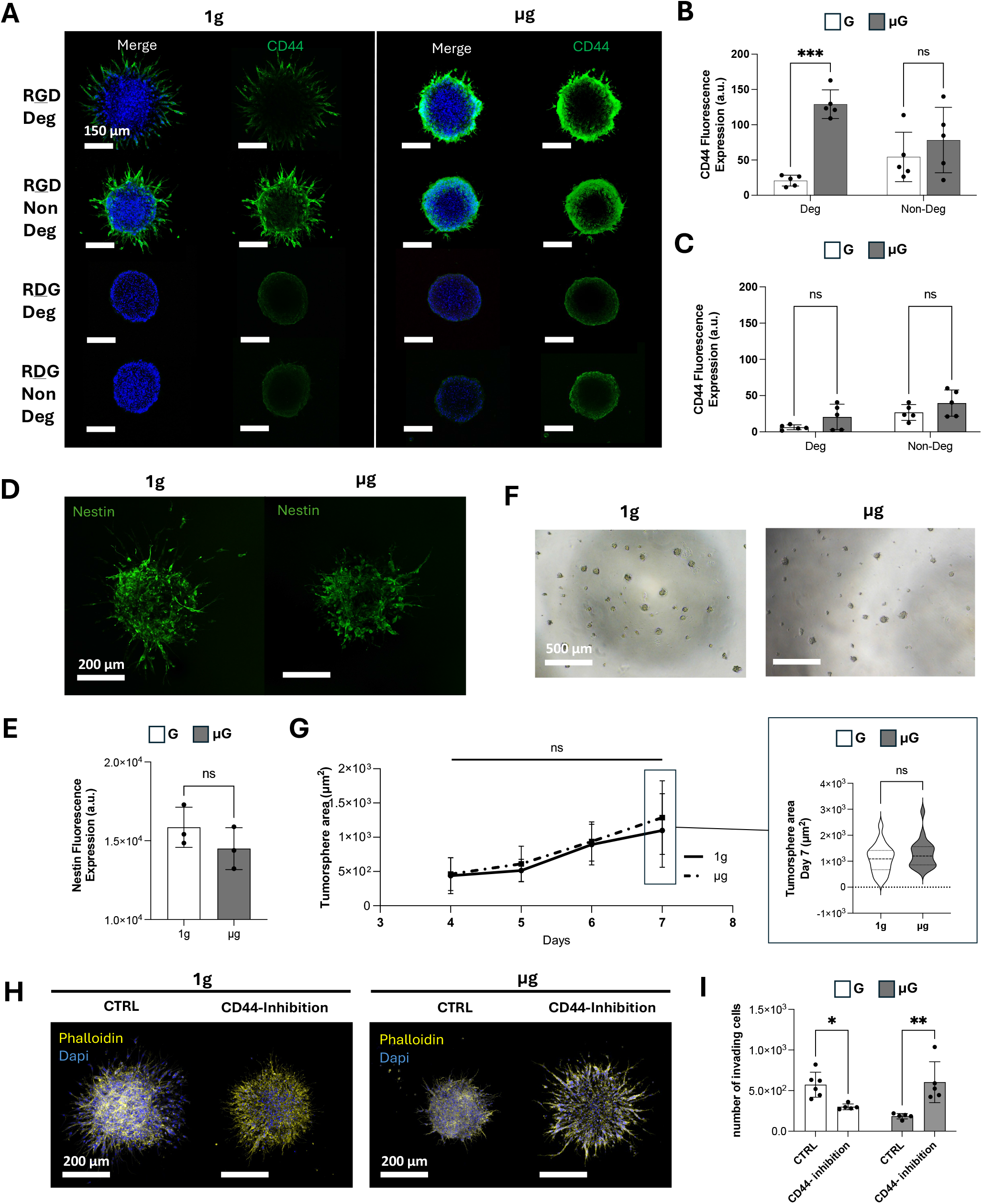
CD44-dependent adhesion response under simulated microgravity and assessment of stemness markers. **(A)** Representative immunofluorescence images of U87 spheroids embedded in adhesive NorHA hydrogels (medium stiffness) and cultured for 24 h under normal gravity (1g) or simulated microgravity (μg). Spheroids were stained with phalloidin (CD44, green) and DAPI (nuclei, blue). Scale bars: 150 μm. **(B)** Quantification of CD44 fluorescence intensity in spheroids cultured in adhesive matrices (R<u>G</u>D/Deg Versus NonDeg) under 1g and μg conditions and expressed as the mean ± SEM from 5 independent experiments. **(C**) Quantification of CD44 fluorescence intensity measured across non-adhesive matrices (R<u>D</u>G/Deg Versus NonDeg) and expressed as the mean ± SEM from 5 independent experiments. **(D)** Representative immunofluorescence images of Nestin immunostaining (green) in spheroids cultured under 1g and μg conditions. Scale bars: 200 μm. **(E)** Quantification of Nestin fluorescence intensity showing no significant difference between gravity conditions. Data are expressed as the mean ± SEM from 3 independent experiments. **(F)** Representative brightfield images of tumorspheres formed from cells recovered after 24 h culture under 1g or μg conditions. Scale bars: 500 μm. **(G)** Quantification of tumorsphere formation efficiency and size over 7 days of culture in serum free media. Inset shows the distribution of tumorsphere area at day 7, represented as a violin plot. (**H)** Representative immunofluorescence images of phalloidin staining (yellow) and dapi (blue) of spheroids cultured under 1g and μg conditions, treated or not treated with CD44 inhibition. **(I)** Quantification of cell dissemination following CD44 inhibition under gravity and simulated microgravity in HA-based matrices and expressed as the mean ± SEM from 5 independent experiments.

We then asked whether this adaptive program extended to canonical stemness pathways. Immunofluorescence staining of Nestin, a progenitor marker often associated with GBM self-renewal, revealed no change in expression under microgravity compared to 1g **(Fig. 4D, E)**. Consistent with this, tumorsphere assays showed no increase in sphere-forming capacity **(Fig. 4F, G)**, indicating that CD44 upregulation under simulated microgravity reflects enhanced adhesion and hyaluronan engagement rather than activation of a stem cell–like program. Functionally, CD44 inhibition decreased cell dissemination under normal gravity, whereas increased dissemination under simulated microgravity **(Fig. 4H, I)**, demonstrating a context-dependent role for CD44.

In mechanically unloaded environments, CD44-mediated adhesion appears to reinforce an over-anchored state that limits cell migration.

### β1-integrin and N-cadherin mediate adhesion-dependent confinement of GBM cells under simulated microgravity

The marked upregulation of CD44 under simulated microgravity suggests a compensatory response aimed at reinforcing cell–matrix interactions through hyaluronan binding. This behavior mirrors our observations in more physically restrictive matrix environments (Fig. 2), where cells similarly increased CD44 expression when motility was limited. Consistent with this, proteomic profiling revealed enrichment of adhesion- and integrin-related signaling pathways under simulated microgravity. We reasoned that, in microgravity, this response could arise from excessive stabilization of adhesion anchors, which limits adhesion dynamics and prevents efficient forward migration. To test this, we next inhibited α5β1-integrin under simulated microgravity in spheroids growth in adhesive and degradable matrices. Remarkably, blocking α5β1-integrin restored cell dissemination from GBM spheroids under unloading **(Fig. 5A, B)**, confirming that excessive β1-mediated adhesion contributes to migration arrest. In contrast, under normal gravity, the same inhibition reduced invasion, consistent with the established pro-migratory role of β1-integrin under standard mechanical conditions. Additionally, inhibiting αvβ3 had no effect under simulated microgravity condition **(Fig. 5C, D)**, indicating the likelihood of selective β1-dependent over-anchorage. Notably, CD44 expression decreased upon β3 integrin inhibition under simulated microgravity but remained largely unchanged in 1g controls **(Fig. 5 E)**, confirming that CD44 upregulation depends on active integrin signaling.

**Fig. 5.**
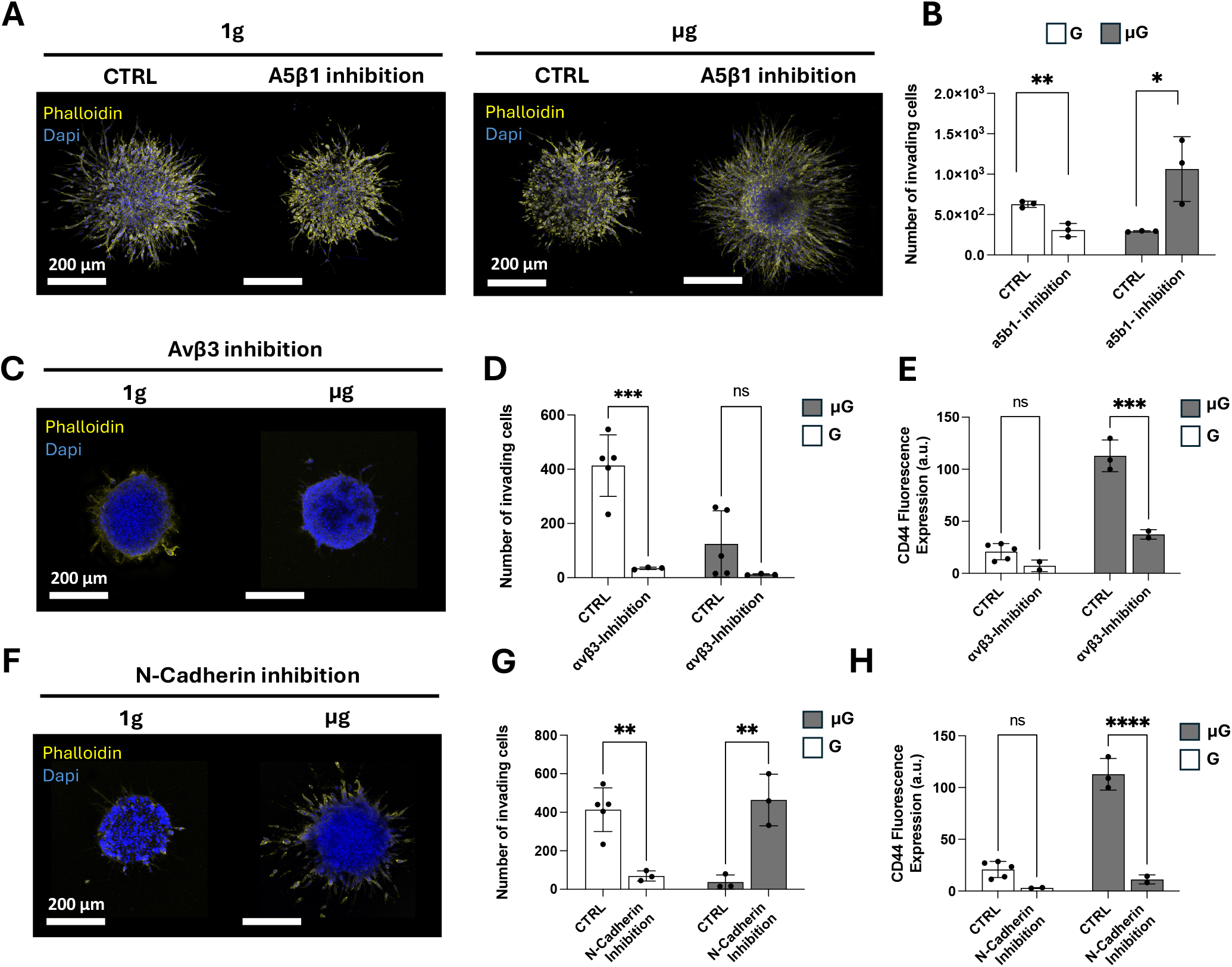
Integrin- and CD44-dependent rewiring of invasion under simulated microgravity. **(A)** Representative immunofluorescence images of U87 spheroids embedded in adhesive, degradable NorHA hydrogels and cultured for 24 h under normal gravity (1g) or simulated microgravity (μg), treated or not with β1-integrin inhibition. Cells were stained for F-actin (phalloidin, yellow) and nuclei (DAPI, blue). Scale bars: 200 μm. **(B)** Quantification of cell invading under 1g and μg conditions following β1-integrin inhibition and expressed as the mean ± SEM from 3 independent experiments. **(C)** Representative immunofluorescence images of U87 spheroids cultured under 1g or μg conditions, treated or not with β3-integrin inhibition. Cells are stained for F-actin (yellow) and nuclei (blue). Scale bars: 200 μm. (D) Quantification of cell invading under 1g and μg conditions following β3-integrin inhibition and expressed as the mean ± SEM from 3 independent experiments, except for relative control (n=5). **(E)** Quantification of CD44 fluorescence intensity following β3-integrin inhibition under 1g and μg conditions and expressed as the mean ± SEM from 2 independent experiments, except for relative control (1g, n=5; μg, n=3). **(F)** Representative immunofluorescence images of spheroids cultured under 1g or μg conditions treated or not with N-cadherin inhibition, stained for F-actin (yellow) and nuclei (blue). Scale bars: 200 μm. **(G)** Quantification of cell invading under 1g and μg conditions following N-cadherin inhibition and expressed as the mean ± SEM from at least 3 independent experiments. **(H)** Quantification of CD44 fluorescence intensity following N-cadherin inhibition and expressed as the mean ± SEM from 2 independent experiments, except for relative control (1g, n=5; μg, n=3).

We next asked whether cell–cell adhesion similarly contributed to the cohesive, migration-restricted phenotype observed under simulated microgravity. Similar to our integrin blocking, inhibiting N-cadherin also restored invasion, reversing the rounded, clustered morphology and re-establishing protrusive mesenchymal-like dissemination. In contrast, under 1g conditions the same treatment reduced invasion, indicating that N-cadherin supports efficient migration when mechanical cues are present but becomes restrictive in a mechanically unloaded environment (**Fig. 5F, G)**. Importantly, N-cadherin blockade under microgravity also reduced CD44 expression **(Fig. 5H)**, linking the microgravity cohesion program directly to the integrin–CD44 axis.

Together, these results indicate that microgravity does not eliminate motility capacity but instead traps GBM cells in an adhesion-dominated state in which excessive engagement of cell–matrix and cell–cell adhesion systems stabilise a cohesive, migration-restricted phenotype.

## Discussion

Glioblastoma (GBM) remains lethal in large part due to its plasticity, which enables tumor cells to rewire invasion strategies and evade therapy. Here, we leverage simulated microgravity as a biophysical perturbation to explore how GBM cells reorganize matrix-mediated invasion when directional mechanical cues are removed, thereby accessing mechanical conditions that cannot be readily reproduced under standard gravity.

Although microgravity has long been recognized as a powerful modulator of cellular mechanosensing^21, 36, 37^, its molecular interpretation has remained constrained by experimental models lacking defined extracellular matrix (ECM) environments. In such systems, poorly defined matrices preclude the establishment of clear matrix structure–cell function relationships, making it difficult to disentangle genuine mechanosensory adaptations from culture-induced artefacts. Recent work has begun to document how brain tumor cells respond under real or simulated microgravity^38^, consistently reporting disruption of cell migration and adhesion, alteration in cytoskeletal organization, and reduced invasive behaviour.^35, 39-43^ Moreover, *Singh et al*. reported that simulated microgravity induces DNA damage and impairs repair pathways in both glioblastoma and microglial cells, highlighting gravitational unloading as a broad cellular stressor affecting survival and genomic integrity.^44^ While painting a striking picture of brain cancer adaptation to microgravity, mechanistic details are still missing, preventing a clear mechanistic understanding of how microgravity influences tumor invasion behavior. This limitation largely reflects the continued reliance on two-dimensional or poorly defined culture systems, in which ECM physicochemical properties cannot be independently controlled, despite their central role in regulating GBM invasion and plasticity. To solve this problem, we established a brain-relevant 3D hyaluronan-based reductionist model, providing versatile control over microenvironmental parameters, thereby reproducing canonical ECM-dependent invasion behaviors. By independently tuning adhesiveness, degradability, and stiffness, the platform recapitulated well-established features of GBM mechanobiology, including the requirement for integrin engagement and matrix degradation to support invasion, and the emergence of confinement-associated phenotypes. With control over matrix-dependent behavior, and simulated microgravity serving as defined mechanical perturbation, we established direct relationships between microenvironmental context and how directional mechanical input reshapes invasion decisions.

Within these controlled systems, removal of gravitational input revealed a marked shift in invasion behavior. Under microgravity conditions, morphological analysis revealed reduced protrusive networks and increased cell-cell and cell-matrix adhesion, accompanied by reduced cell proliferative activity, consistent with previous reports showing reduced tumor cell growth, metabolism, and invasiveness in three-dimensional scaffold models.^45^ Various “Omics” analyses have reported differential regulation of ECM remodeling pathways under microgravity, including reduced proteolytic activity, decreased integrin signaling, perturbation in cell cycle regulators, and increased prevalence of apoptosis.^46-48^ However, these observations are typically not resolved within a defined microenvironmental framework. Using our system, proteomic profiling revealed enrichment of cell–cell junction assembly and adhesion-related processes alongside downregulation of canonical motility regulators, providing a mechanistic link between these molecular changes and the emergence of an adhesion-dominated invasion state. Consistent with this interpretation, simulated microgravity induced junctional localization of N-cadherin and cortical redistribution of phosphorylated myosin light chain, both indicative of increased cell–cell adhesion and cortical contractility^49, 50^. Together, these features suggest a shift toward reinforced adhesion and altered force transmission under mechanical unloading. In line with this, recent work from Poole and colleagues demonstrated that simulated microgravity alters focal adhesion organization, mechanosensitive signaling, and 3D invasion in an ELKIN1-dependent manner, highlighting that gravitational unloading directly perturbs force-sensing pathways^31^.

Given this shift toward an adhesion-dominated state, we next examined whether key adhesion receptors associated with GBM invasion were similarly modulated. CD44, a principal hyaluronan receptor, is widely implicated in GBM invasion, stemness, and tumor progression, and is often associated with increased tumor aggressiveness and dissemination. Strikingly, we observed robust upregulation of CD44 under microgravity despite an overall reduction in invasion. However, this increase was not accompanied by broader evidence of enhanced stemness, as neither Nestin expression nor tumoursphere formation was increased, which contrasts with previous reports linking elevated CD44 expression to enhanced invasion and tumourigenicity.^8, 51^ Therefore, we speculate that increased CD44 in this context relates to the observed adhesion phenotype. Importantly, this increase was specific to hyaluronan-based matrices and was not observed in a general polysaccharide control (alginate), which lack CD44-binding capacity, confirming the importance of hyaluronan in regulating this phenotype. Targeted inhibition of CD44 under normal gravity reduced invasion, consistent with its role in supporting mesenchymal, adhesion-driven migration,^52^ whereas under simulated microgravity the same inhibition restored cell dissemination. This suggests that, under mechanical unloading, CD44-mediated adhesion contributes to retaining cells within a highly anchored state, limiting their ability to migrate. Rather than acting as a pro-invasive marker, CD44 displays a context-dependent role, supporting migration under normal gravity but contributing to an adhesion-dominated state that limits dissemination under mechanical unloading.

The CD44-dependent adhesion shift observed under unloading, together with its restriction to adhesive (RGD-modified) matrices, suggests that integrin engagement contributes to stabilizing this adaptive state. Consistent with the proteomic enrichment of adhesion- and integrin-associated pathways, functional perturbation revealed that β1-integrin plays a central role in constraining invasion under simulated microgravity. Inhibition of β1-integrin restored invasion under unloading, whereas β3 inhibition did not, indicating β1-dependent over-anchorage rather than a general impairment of motility. A similar context dependence was observed for cell– cell adhesion: N-cadherin inhibition restored invasion under microgravity but reduced invasion under normal gravity. This indicates that adhesion signaling is not intrinsically pro-invasive but instead operates in a mechanically defined context in which its function can shift from permissive to restrictive. Moreover, β1 inhibition did not alter CD44 levels (data not shown), suggesting that CD44 upregulation is not simply a downstream consequence of β1-mediated adhesion. In contrast, β3 inhibition significantly reduced CD44 expression specifically under microgravity, despite leaving invasion low, implying that β3 signaling contributes to sustaining the CD44-high adaptive program without directly mediating the invasion break. Mechanistically, this decoupling is consistent with integrin subtype specialization in signalling.^53^ Under unloading, β1 appears to control the physical adhesion dynamics that govern mesenchymal migration, whereas β3 activity may have migration independent roles like transcription regulation.

In parallel with this adhesion-dominated phenotype, proteomic analysis of conditioned media together with direct particle tracking revealed increased extracellular vesicle secretion under microgravity, without changes in vesicle size distribution. This suggests that, under microgravity, GBM cells become less physically invasive but remain biologically active, shifting from migration-based dissemination toward communication-based adaptation. In this context, when physical dissemination is constrained, cells may partially compensate by enhancing paracrine communication rather than migratory spread, consistent with recent observations in breast cancer under simulated microgravity.^50^

Together, these findings support a model in which simulated microgravity does not globally suppress GBM motility but instead reconfigures invasion in a ECM-dependent manner. Under permissive conditions, where integrin engagement and matrix degradation normally sustain mesenchymal dissemination, gravitational unloading shifts cells toward an adhesion-dominated state characterized by elevated β1-integrin, CD44 and N-cadherin. Notably, this phenotype closely parallels the behavior observed in physically confining matrix environments at normal gravity condition, such as high-density or non-degradable hydrogels, where invasion is similarly restricted and CD44 expression is increased. Additionally, this behavior aligns with the broader concept that cell migration depends on an optimal balance of adhesion strength, where insufficient adhesion prevents traction, while excessive adhesion becomes restrictive^53-58^. In this context, simulated microgravity can be interpreted not as a creating a new phenotype, but as shifting GBM cells beyond a permissive adhesion regime toward a confinement-like state in which reinforced HA–CD44 engagement, β1-integrin anchoring, and N-cadherin-mediated cohesion collectively restrict dissemination.

## Conclusion

In this work, we combined simulated microgravity with a tunable NorHA–PEG hydrogel platform to dissect how GBM adapts invasion when directional mechanical cues are disrupted. By independently controlling adhesiveness, degradability, and stiffness, the system provided a defined three-dimensional reference state in which microgravity could be applied as a physical perturbation. Under these conditions, gravitational unloading suppressed mesenchymal dissemination and shifted cells toward an adhesion- and cohesion-dominated state involving HA–CD44 engagement, β1-integrin signaling, and N-cadherin. Notably, adhesion pathways that normally promote invasion became restrictive under microgravity, revealing a context-dependent mechanosensitive vulnerability. More broadly, these findings demonstrate how combining ECM engineering with controlled mechanical perturbation access to invasion states that are otherwise difficult to isolate, providing a framework to study how physical cues regulate cancer cell behavior.

## Author contributions

G.S. and K.A.K. conceived the study and initiated the project. G.S. designed and performed experiments across the full range of approaches, analyzed the data, prepared the figures, and coordinated manuscript preparation. C.W. contributed to spheroid encapsulation in 3D matrices and microgravity exposure experiments. N.W. and Y.L. characterized the 3D hydrogel model through its design and rheological analysis. K.A.S. performed the proteomic analysis. R.B.D. performed the extracellular vesicle analysis. M.S. contributed to interpretation and discussion of the results. K.P. provided reagents for the mechanistic studies and contributed to discussion of the findings. K.A.K. supervised all aspects of the work and contributed to experimental design and manuscript organization. All authors contributed to writing and revising the manuscript.

## Supplementary Figures

**Fig. S1.**
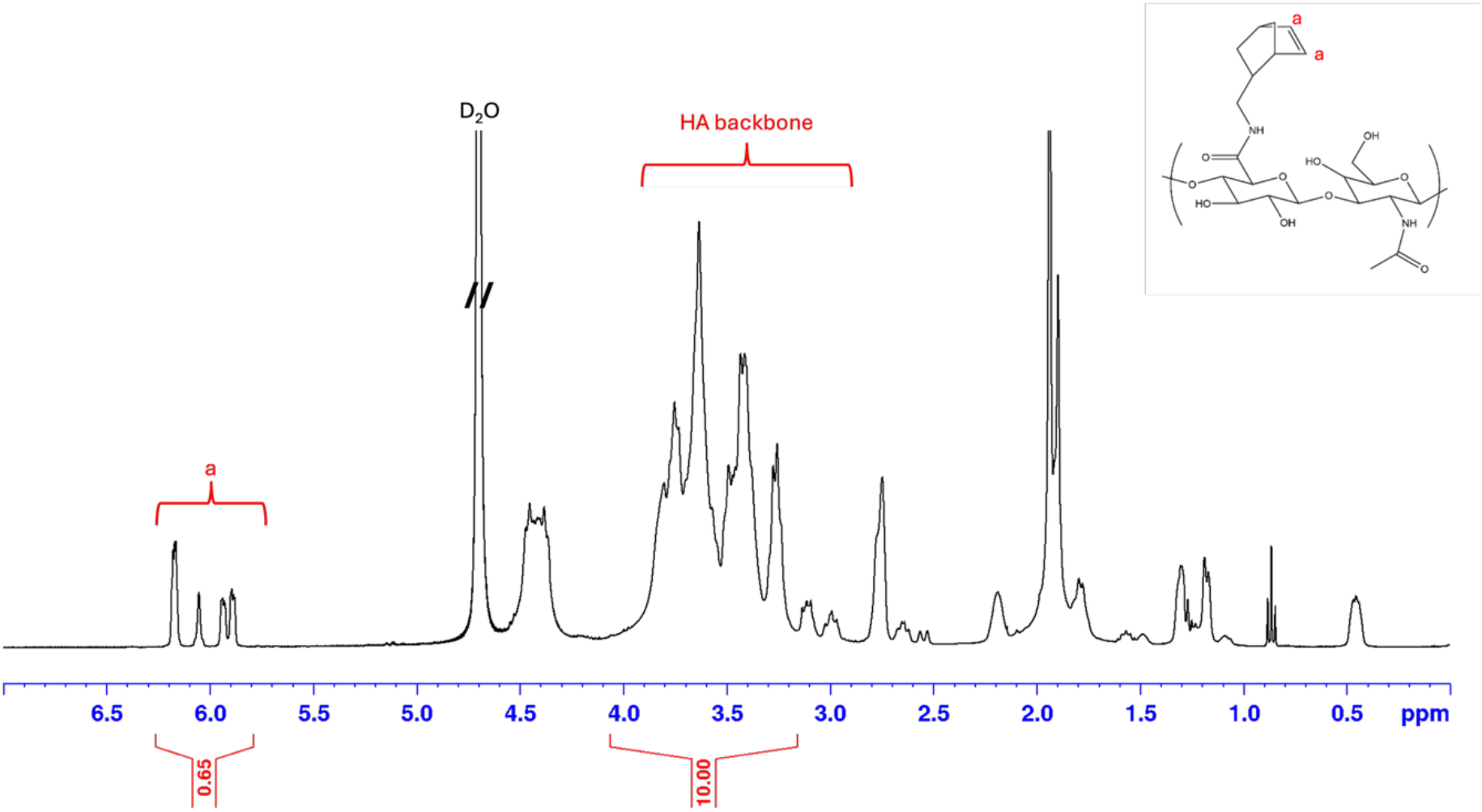
Structural confirmation of norbornene-functionalised hyaluronic acid (NorHA) by ^1^H NMR spectroscopy. Representative ^1^H NMR spectrum of the modified polymer showing the characteristic resonances associated with norbornene functional groups alongside the native proton signals of the hyaluronic acid backbone. The appearance of distinct olefinic signals in the downfield region confirms successful incorporation of the norbornene moieties, while the broad multiplet region corresponds to protons from the polysaccharide backbone. The schematic structure of the modified repeat unit is shown for reference, illustrating the chemical modification of hyaluronic acid with norbornene groups.

**Fig. S2.**
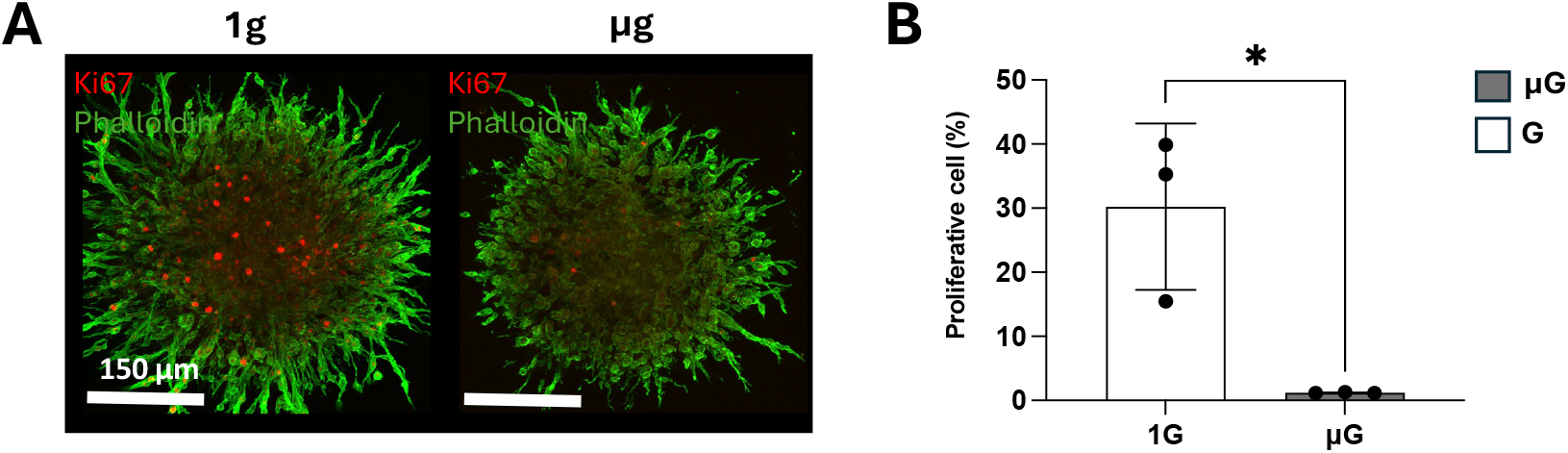
Simulated microgravity reduces proliferative activity in GBM spheroids. **(A)** Representative immunofluorescence images of U87 spheroids cultured in adhesive and degradable NorHA/PEG (medium stiffness) hydrogels under normal gravity and simulated microgravity conditions. F-actin cytoskeleton is shown by phalloidin staining (green), while proliferating cells are identified by Ki67 staining (red). **(B)** Quantification of Ki67-positive cells per spheroid confirming a reduction in proliferative activity under simulated microgravity conditions. Scale bar: 150 µm. Data are expressed as the mean ± SEM from 3 independent experiments.

**Fig. S3.**
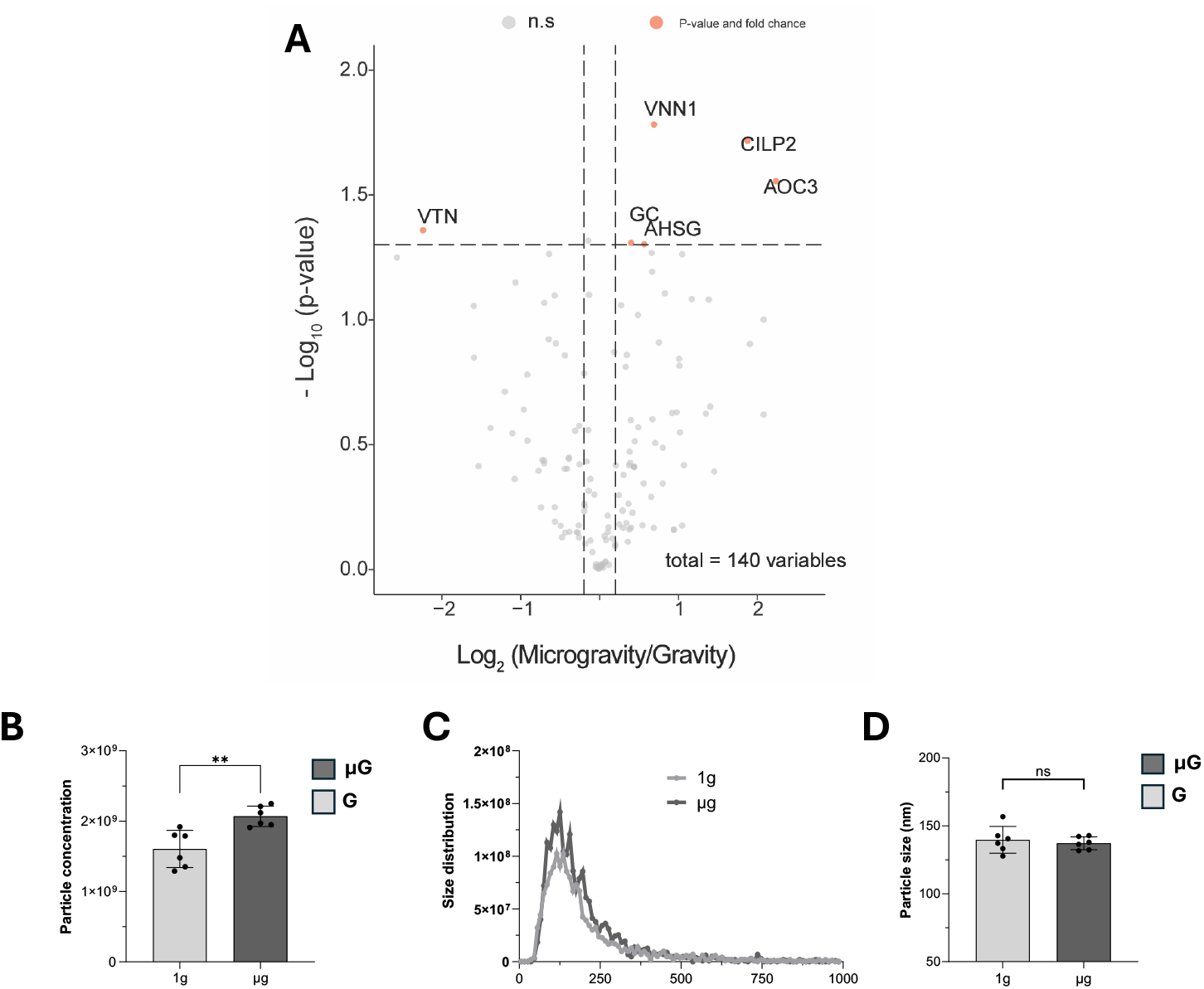
Proteomic profiling reveals microgravity-associated changes in vesicle-related proteins. **(A)** Volcano plot showing differential protein abundance in conditioned media collected from GBM spheroids cultured under simulated microgravity compared with normal gravity conditions. The x-axis represents the log_2_ fold change (microgravity/gravity), and the y-axis shows the –log_10_(p-value). Dashed lines indicate statistical and fold-change thresholds used to identify differentially expressed proteins. Proteins significantly upregulated under simulated microgravity are highlighted in orange. **(B)** Quantification of extracellular vesicles (EVs) from spheroids cultured under simulated microgravity compared with normal gravity controls. **(C)** Size distribution of EVs showing no difference between the two groups. **(D)** Quantification of EVs size from spheroids cultured under simulated microgravity and normal gravity. Data are presented as mean ± S.D. from 6 independent experiments.

**Fig. S4.**
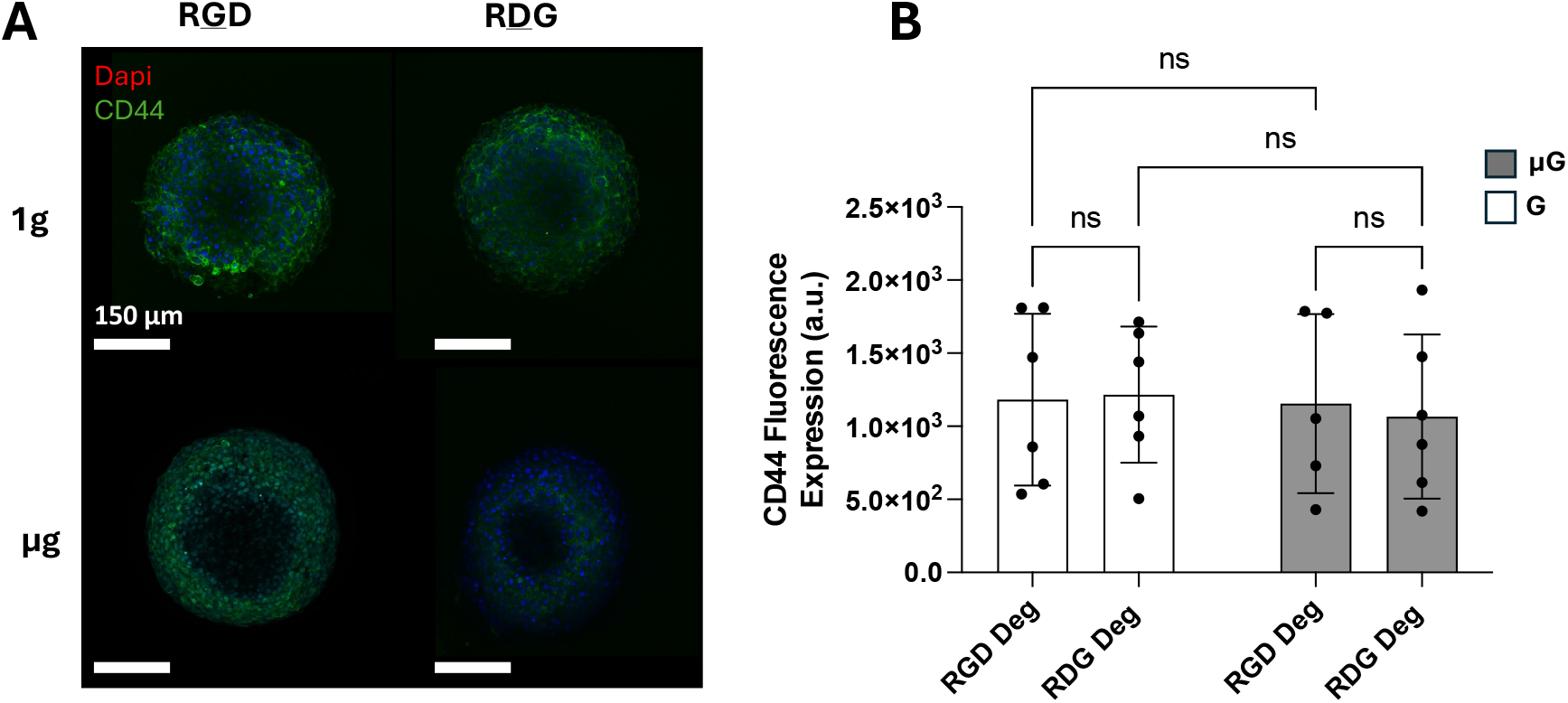
CD44 upregulation under simulated microgravity requires hyaluronan engagement and is not reproduced in alginate matrices. **(A)** Representative immunofluorescence images of GBM spheroids cultured in alginate hydrogels under normal gravity and simulated microgravity conditions. CD44 is shown in green and nuclei are stained with DAPI (blue). Alginate hydrogels were used in both non-adhesive and adhesive (RGD-modified) formulations to control for integrin-mediated cell–matrix adhesion. Scale bar: 150 µm. **(B)** Quantification of CD44 fluorescence intensity in spheroids cultured under the indicated conditions. No significant differences in CD44 expression were observed between normal gravity and simulated microgravity in either adhesive or non-adhesive alginate matrices, indicating that integrin-mediated adhesion alone does not drive CD44 upregulation. Data are presented as mean ± S.D. from 6 independent experiments.

